# Using Attention-based Deep Learning to Predict ERG:TMPRSS2 Fusion Status in Prostate Cancer from Whole Slide Images

**DOI:** 10.1101/2022.11.18.517111

**Authors:** Mohamed Omar, Zhuoran Xu, Sophie B Rand, Mohammad Mohammad, Daniela C. Salles, Edward M. Schaeffer, Brian D. Robinson, Tamara L. Lotan, Massimo Loda, Luigi Marchionni

## Abstract

Prostate cancer (PCa) is associated with several genetic alterations which play an important role in the disease heterogeneity and clinical outcome including gene fusion between TMPRSS2 and members of the ETS family of transcription factors specially ERG. The expanding wealth of pathology whole slide images (WSIs) and the increasing adoption of deep learning (DL) approaches offer a unique opportunity for pathologists to streamline the detection of ERG:TMPRSS2 fusion status. Here, we used two large cohorts of digitized H&E-stained slides from radical prostatectomy specimens to train and evaluate a DL system capable of detecting the ERG fusion status and also detecting tissue regions of high diagnostic and prognostic relevance. Slides from the PCa TCGA dataset were split into training (n=318), validation (n=59), and testing sets (n=59) with the training and validation sets being used for training the model and optimizing its hyperparameters, respectively while the testing set was used for evaluating the performance. Additionally, we used an internal testing cohort consisting of 314 WSIs for independent assessment of the model’s performance. The ERG prediction model achieved an Area Under the Receiver Operating Characteristic curve (AUC) of 0.72 and 0.73 in the TCGA testing set and the internal testing cohort, respectively. In addition to slide-level classification, we also identified highly attended patches for the cases predicted as either ERG-positive or negative which had distinct morphological features associated with ERG status. We subsequently characterized the cellular composition of these patches using HoVer-Net model trained on the PanNuke dataset to segment and classify the nuclei into five main categories. Notably, a high ratio of neoplastic cells in the highly-attended regions was significantly associated with shorter overall and progression-free survival while high ratios of immune, stromal and stromal to neoplastic cells were all associated with longer overall and metastases-free survival. Our work highlights the utility of deploying deep learning systems on digitized histopathology slides to predict key molecular alteration in cancer together with their associated morphological features which would streamline the diagnostic process.

## Introduction

Prostate cancer (PCa) is the second most common solid organ malignancy among men worldwide with more than 1 million annually diagnosed new cases ^1,2^. The PCa genome is characterized by several gene rearrangements involving E26 transformation-specific (ETS) transcription factors ^3^. Of the ETS rearrangements, fusion between ERG and the androgen-regulated transmembrane protease, serine 2 (TMPRSS2) is the most frequent, being present in nearly 40-50% of PCa cases ^4^. The ERG:TMPRSS2 fusion was found to be instrumental during the transformation of prostate intraepithelial neoplastic lesions into invasive adenocarcinoma as well as during PCa progression and metastasis ^5^.

The diagnosis of PCa is based on the light microscopic examination of hematoxylin and eosin (H&E)-stained tissue sections obtained from the prostate gland ^6^–8. However, H&E staining alone cannot differentiate between ERG-positive (those with ERG:TMPRSS2 fusion) and ERG-negative tumors. Instead, ERG status is usually detected using fluorescence in situ hybridization (FISH) or reverse transcription-polymerase chain reaction (RT-PCR) while immunohistochemical staining for the ERG protein can be used to infer the ERG:TMPRSS2 gene fusion status with optimal sensitivity and specificity ^9^. Since these technologies are costly and require specialized equipment and trained personnel, there is need for innovative tools to decrease the cost and streamline the diagnostic process. Specifically, a tool that can utilize routine H&E-stained slides for such task would potentially achieve the aforementioned goals given the practicality and widespread availability of these slides.

Since 2000, whole-slide imaging (WSI) started to become common, as digital slide scanners became commercially available ^10^. Currently, modern pathology practice is moving toward a digital workflow through the deployment of artificial intelligence (AI) and computer vision systems for image preprocessing, segmentation, feature detection, and quantification on H&E-stained WSIs ^10,11^. Deep learning (DL) systems in particular have been employed extensively over the past few years in several tasks involving the use of H&E-stained WSIs for phenotype prediction, classification, and subtyping especially in cancer research ^12,13^. For instance, DL models have been developed to automatically detect tumor regions in several cancer types including breast, lung, and prostate cancers ^14–19^. Additionally, DL has been employed in more advanced tasks including classification of tumor subtypes ^20–22^, tumor grading ^19,23^, and prediction of therapeutic response ^24^. Notably, the utilization of DL systems in molecular pathology has gained momentum over the past years with more models being deployed to predict genetic alterations from histopathology images ^12^. For instance, Coudray et al. have developed a model for predicting several key mutations in lung adenocarcinoma ^25^ while Bilal et al. developed a DL system for predicting several key genetic alterations in colorectal cancer ^26^. Similar studies have aimed at identifying other molecular alterations and phenotypes including ER status in breast cancer ^27^, BRAF mutations in melanoma ^28^, and SPOP mutations in prostate cancer ^29^.

Here, we introduce a semi-supervised DL model capable of predicting the ERG:TMPRSS2 gene fusion status solely from H&E-stained pathology images. This model has been trained and validated using two large imaging cohorts containing 750 WSIs derived from radical prostatectomy specimens of PCa patients with known ERG status. Additionally, we deciphered the cellular composition of the top patches contributing to the prediction of each ERG phenotype and examined the association of this cellular composition with overall, progression-free, and metastasis-free survival.

## Materials & Methods

### Patients and Slides Selection

We queried the Genomic Data Commons (GDC) portal and the Cancer Genome Atlas (TCGA) for whole slide images (WSIs) derived from primary prostate cancer (PCa) patients with known TMPRSS2:ERG gene fusion status. The following terms were used: Project Id: TCGA-PRAD, Data Type: Slide Image, Experimental Strategy: Diagnostic Slide. The PCa TCGA dataset included 436 formalin-fixed paraffin-embedded (FFPE) H&E-stained WSIs from 393 PCa patients with known ERG fusion status (positive versus negative) determined by FusionSeq ^3,30^. Subsequently, a dataset of 314 FFPE-derived WSIs from PCa patients has been provided by the Johns Hopkins University (henceforth termed the natural history cohort) and was used as an additional testing set. The ERG status in this cohort has been determined using IHC staining of FFPE tumor tissue from radical prostatectomy specimens to detect the ERG/TMPRSS2 fusion protein ^9^. Slides from the TCGA and natural history cohorts were digitized using Aperio (Leica Biosystems) and Hamamatsu (Hamamatsu Photonics K.K.) slide scanners, respectively.

### Image Preprocessing

#### Tissue Segmentation

Tissue segmentation was performed on all slides by first converting the RGB images to HSV space then applying a median blur filter with a kernel size of 7, followed by thresholding using Otsu’s method ^31^. Finally, morphological closing (dilation followed by erosion) with a 4×4 structuring element was applied to close small gaps.

#### Tiling and Feature Extraction

Following segmentation, tissue masks were tiled into 2048×2048 patches at ×40 magnification. We followed by down sampling the patches by a factor of 4 resulting into 512×512 patches at ×10 magnification on which feature extraction was performed using a pre-trained ResNet50 model ^32^.

### Training the ERG Status Model

#### The Deep Learning Framework

To build a model capable of detecting the ERG status from H&E WSIs, we used Clustering-constrained Attention Multiple Instance Learning (CLAM) ^32^. CLAM is a modified multiple instance learning (MIL) framework which aggregates patch-level into slide-level representations using attention-based pooling function instead of max pooling ^33^. In the CLAM network, the first layer is a fully connected linear layer that takes the 1024-dimensional vector representing the extracted patch features and returns a 512-dimensional vector which is then fed to the attention network. The attention network is based on a gated attention mechanism that assigns different weights to instances (patches) within a bag (WSI) based on their contributions to the slide-level prediction ^33^. This network then splits into 2 separate branches, one for each class (ERG-positive and negative). Notably, slide-level representations are scored by 2 class-specific separate classifiers and a softmax function is then used to convert these into class-specific probability scores for each WSI. Cross-entropy loss was used to compare the slide-level predictions to the true labels and the model’s weights were modified using Adam Optimizer with an alpha of 0.0001 and weight decay of 0.00001. Finally, a maximum of 150 epochs was used for training the model and we used early stopping to stop training and save the model if the error in the validation set did not decrease for over 20 epochs.

#### WSIs Datasets

We divided the TCGA PCa cohort into training (70%), validation (15%), and testing (15%) sets. The training (n=318 WSIs) and validation (n=59) sets were used for training the model and tuning its hyperparameters, respectively while the testing set (n=59) was left out for evaluating the model’s performance.

### Independent Evaluation of Performance

In addition to the testing set derived from the TCGA cohort (n=59), we tested the model on the entire natural history cohort (n=314) to provide a completely independent assessment of performance. The model was used to infer class-specific predicted probabilities for each WSI in the natural history cohort. Using the best threshold from the ROC curve of the training data, we converted these probability scores into binary class predictions (ERG positive vs negative) and compared these predictions with the true class labels.

### Nuclei Segmentation and Classification

To decipher the cellular composition of the highly attended regions by our model, we used the HoVer-Net model trained on the PanNuke dataset ^34,35^ to segment and classify the nuclei in these regions. Specifically, we pooled the patches with highest attention scores (15 patches per WSI) from all WSIs predicted as either positive or negative. Notably, these patches were retrieved after basic preprocessing and tissue segmentation and had the size of 512×512 at ×10 magnification. HoVer-Net was used on patches from each class separately to segment and classify the nuclei into five different types: benign epithelial, neoplastic, inflammatory/immune, necrotic, and stromal ^34^. Subsequently, we compared the abundance of different nuclear types in the highly attended regions from cases predicted as ERG-positive with those predicted as ERG-negative in both the TCGA and natural history cohorts.

### Survival Analyses

To examine the association between the nuclear/cellular content in the highly attended patches and the survival probability, we calculated the number and ratio of different nuclear types in each WSI in the TCGA and natural history cohorts. The ratio of each nuclear type was computed by dividing the absolute number of each nuclear type by the number of all nuclei in the highly attended patches of each slide (top 15 patches). This ratio was then binarized into high versus low content using maximally selected log-rank statistics to compute the best cutoff that is most significantly associated with the survival outcome ^36,37^. We subsequently computed the association between the binarized nuclear content and the overall survival (OS) (TCGA and natural history cohorts), progression-free survival (PFS) (TCGA cohort), and metastases-free survival (natural history cohort) using Kaplan Meier (KM) survival curves ^38^. Notably, only the correctly classified slides from each cohort were included in this analysis.

### Statistical Analyses and Software

The performance was assessed using the AUC, accuracy, balanced accuracy, sensitivity, specificity, and Matthews Correlation Coefficient (MCC). Receiver Operating Characteristics (ROC) curves were plotted using the predicted probability scores together with the ground truth labels. For all cohorts, the probability scores were binarized into predicted classes using the best threshold from the training data. Slide preprocessing and training the deep learning model were performed using *python* (v3.7.5), *openslide* (v3.4.1), and *PyTorch* (v1.3.1). For nuclei segmentation and classification, we used the pytorch (v1.6) implementation of HoVer-Net model trained on the PanNuke dataset ^34,35^. Survival analyses were performed using the *survival* (v3.3-1) ^39^ and *survminer* (v0.4.9) ^40^ packages.

## Results

### Patient Selection

The PCa TCGA cohort included 436 slides from 393 unique patients with available slide-level information about the ERG fusion status. This cohort was split into 70% training (318 WSIs; 188 ERG-negative and 130 ERG-positive), 15% validation, and 15% testing (59 WSIs; 33 ERG-negative and 26 ERG-positive for each). Additionally, we used the entire natural history cohort as an independent testing set to further validate the model’s performance. This cohort included 314 WSIs (185 ERG-negative and 129 ERG-positive) from patients who underwent radical prostatectomy between 1992 and 2010 and received no treatment prior to the procedure.

### Predicting ERG Status From H&E-stained Whole Slide Images

The ERG prediction model was trained on H&E-stained WSIs to distinguish slides derived from patients with ERG-positive from those with ERG-negative using tissue morphological and spatial features only. Since ERG fusion is known to induce changes in the tumor microenvironment, we hypothesized that this can also be associated with morphological changes that are not limited only to the tumor but also involve the stroma and other regions surrounding the tumor. With this in mind, we used the whole tissue sections (after preprocessing) for making predictions instead of using only tumor regions. We trained 10 different models using 10 different splits of the TCGA dataset, with each split consisting of training (n=318), validation (n=59), and testing (n=59) sets (Figure 2A). The average Area Under the ROC Curve (AUC) for the 10 models in the training data was 0.79 (SD=0.03) while the average accuracy was 0.71 (SD=0.03). The best performing model had an AUC of 0.84 and accuracy of 0.77 in the training data (Figure 2A). We further evaluated this model on the TCGA testing set (n=59), in which it had an AUC of 0.72 and accuracy of 0.70 (Figure 2B).

**Figure 1.**
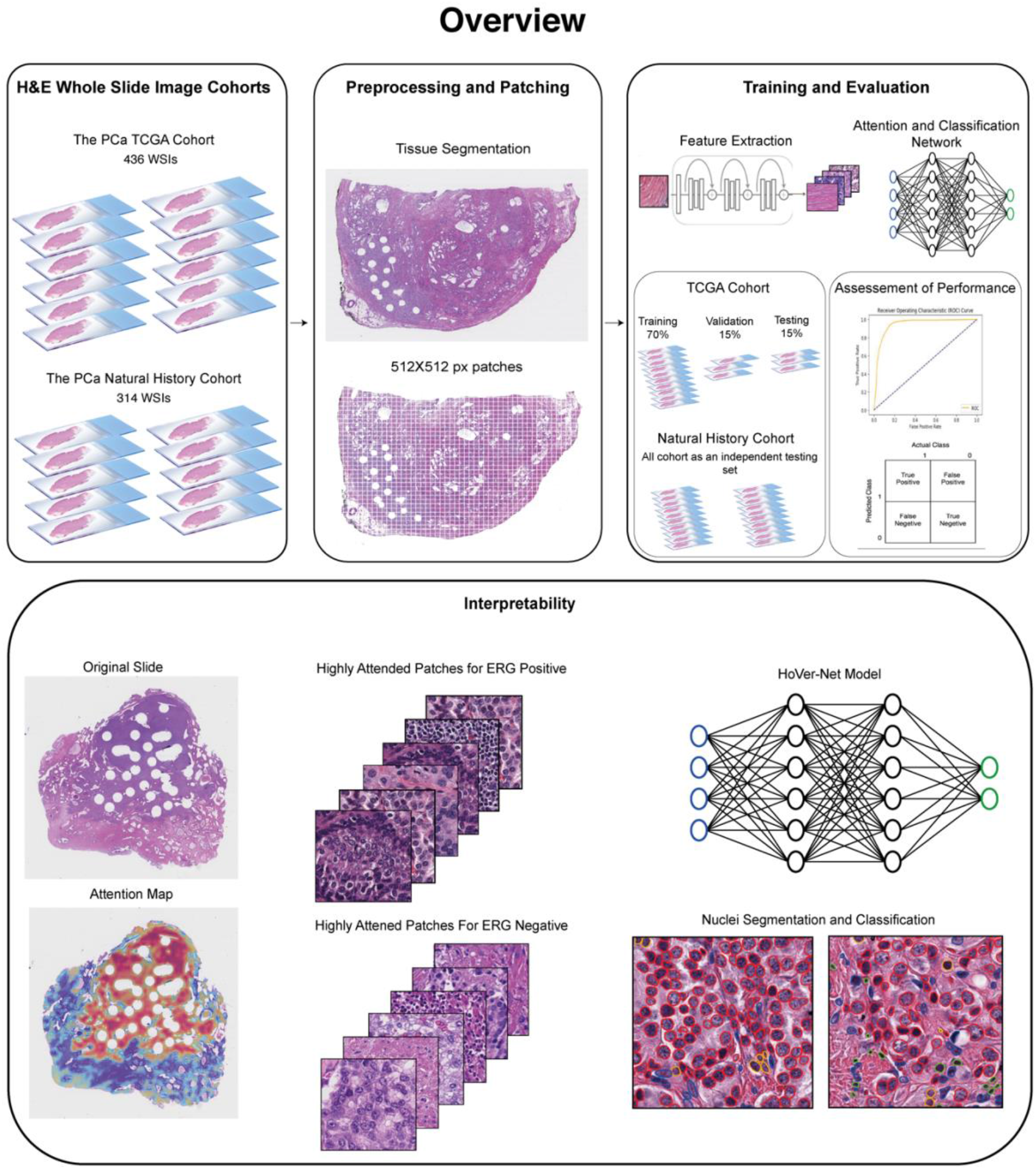
Detecting ERG status using H&E-stained radical prostatectomy specimens from patients with prostate cancer. Whole slide images from the PCa TCGA (n=436) and natural history (n=314) cohorts were used in this study. Following tissue segmentation, each WSI was tiled into 2048×2048 patches at ×40 magnification and were further down sampled by a factor of 4 to extract features from 512×512 patches at ×10 magnification. WSIs from the PCa TCGA cohort were split into training (70%), validation (15%), and testing (15%) sets while the natural history cohort was used as an additional testing set. HoVer-Net model was used for nuclei segmentation and classification in the top patches of WSIs predicted as ERG fusion positive or negative.

**Figure 2.**
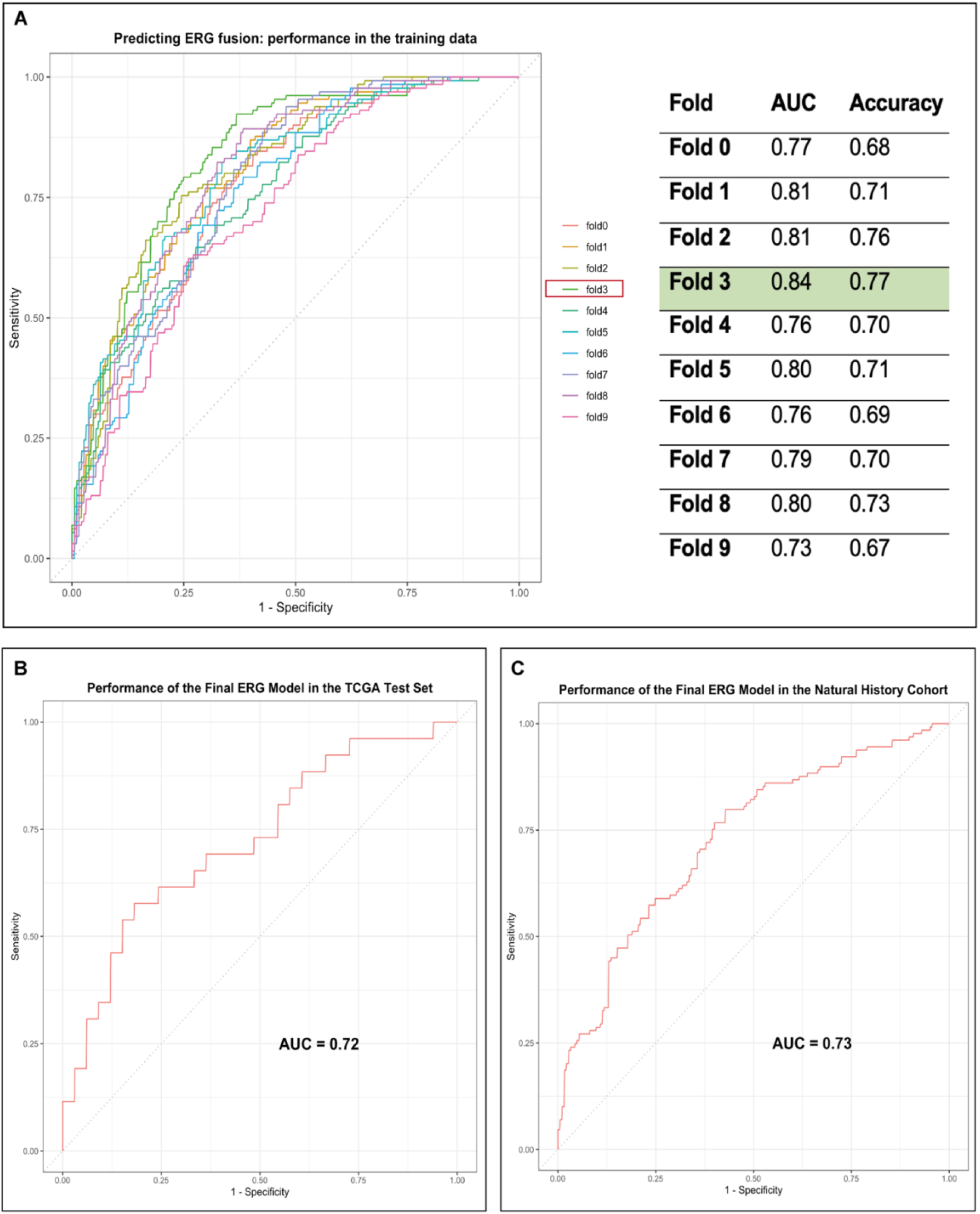
Performance of the models predicting ERG-TMPRSS2 fusion status using H&E-stained whole slide images. **A)** Performance of the models in the 10 different training folds. The prostate cancer TCGA cohort was divided 10 times into training, validation, and testing sets with each fold having different slides in each set. In each fold, models were trained on the training set while the validation set was used for tuning the model hyperparameters. The best performing model (fold 3) was used for downstream evaluation on the TCGA testing set as well as the natural history cohort. **B, C)** Performance of the model in the TCGA testing set (n=59) and the natural history cohort (n=314).

**Figure 3.**
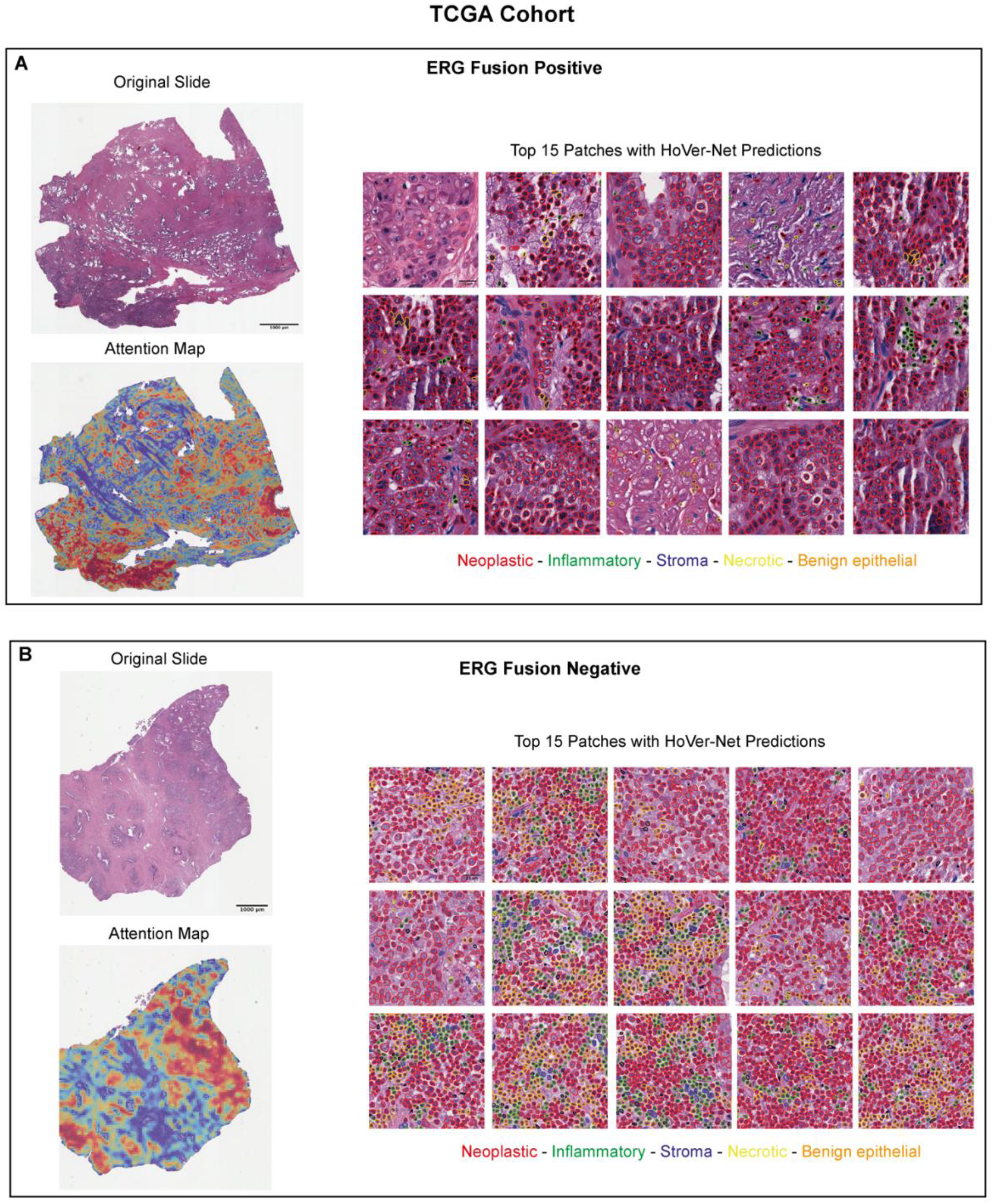
Distinct morphological features corresponding to ERG:TMPRSS2 fusion in the TCGA cohort. **A, B)** Example of a slide predicted as ERG positive (A) and negative (B) with the corresponding top 15 tiles with highest attention scores. HoVer-Net model was used to segment and classify the nuclei in these patches into 5 types: neoplastic, inflammatory, stroma, necrotic, and benign epithelial.

**Figure 4.**
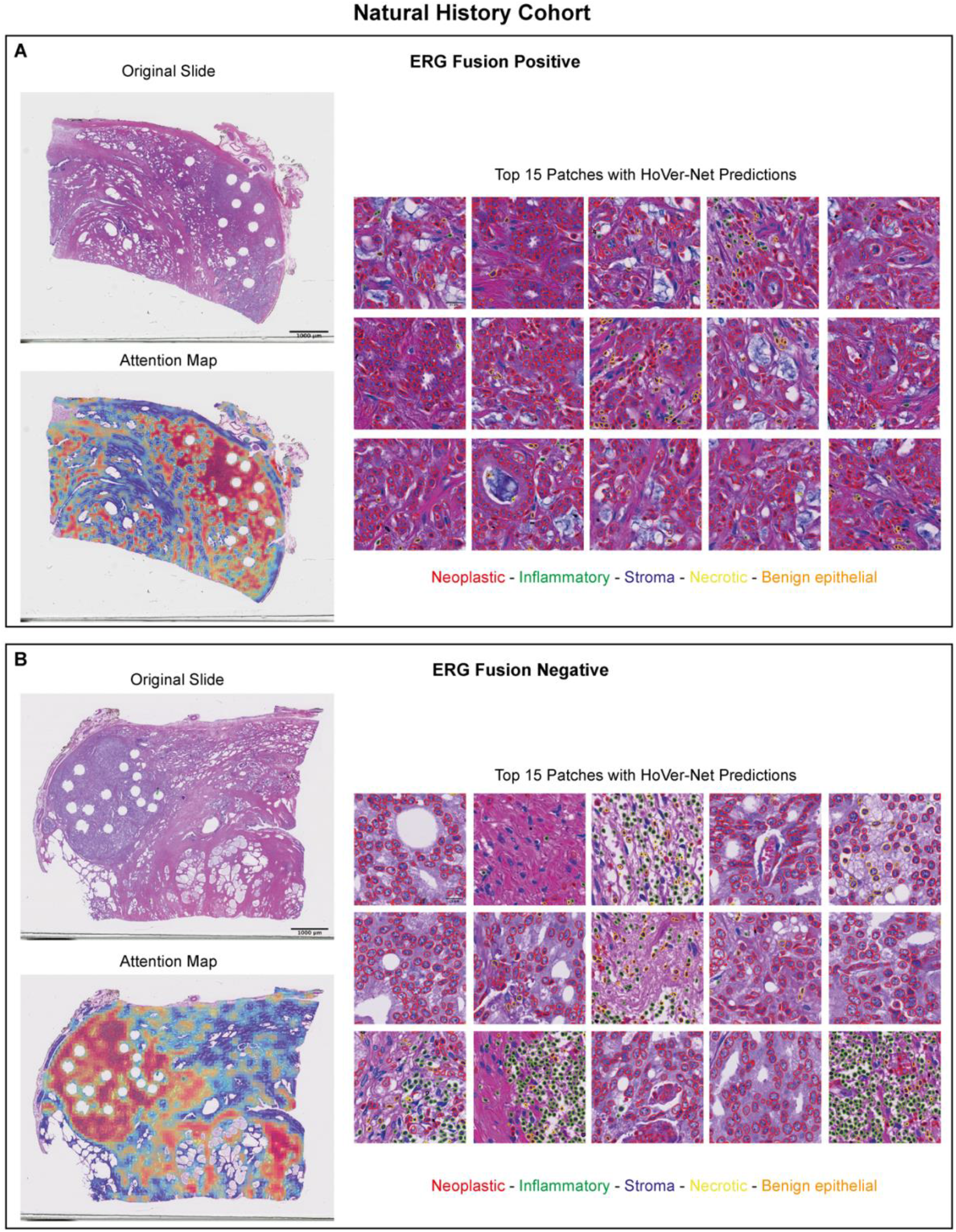
Distinct morphological features corresponding to ERG:TMPRSS2 fusion in the natural history cohort. **A, B)** Example of a slide predicted as ERG positive (A) and negative (B) with the corresponding top 15 tiles with highest attention scores. HoVer-Net model was used to segment and classify the nuclei in these patches into 5 types: neoplastic, inflammatory, stroma, necrotic, and benign epithelial.

### Independent Evaluation of Performance on the Natural History Cohort

To further assess the performance of the ERG fusion status prediction model, we used it to distinguish ERG-positive from negative cases in the natural history cohort which included 314 WSIs of tissue from radical prostatectomy specimens in which the ERG status has been inferred using immunohistochemistry (IHC). In this cohort, the ERG model could detect the ERG status with an AUC of 0.73 and accuracy of 0.69 (Figure 2C).

### Highly Attended Patches Show Distinct Morphological Features Associated with ERG Fusion

To further decipher the cellular architecture contributing to the model’s prediction, we extracted the top 15 highly attended patches (highest attention scores) from each slide predicted as either ERG-positive or negative. We then used HoVer-Net model to perform nuclear segmentation and classification into five categories: benign epithelial, tumor, stroma, inflammatory/immune, and necrotic cells and compared the frequency of these nuclear types between the two predicted classes.

Subsequently, we compared the cellular composition in these top patches between the ERG-positive and negative cases. On average, the highly attended patches from the TCGA WSIs predicted as ERG-positive tended to have more neoplastic content compared to those predicted as negative. The same pattern was observed in the natural history cohort in which the ERG-positive highly attended patches had a higher neoplastic content compared to the ERG-negative patches.

### Cellular Composition in the Highly Attended Patches Is Associated with Survival

We further examined whether the cellular composition in the highly attended patches is associated with the survival probability in both cohorts. For each WSI correctly predicted as positive or negative, we computed the number and ratio of each nuclear type predicted by the HoVer-Net model in the highly attended patches (top 15 patches with the highest attention scores for each WSI). We subsequently computed the association between the cellular composition and the progression-free survival in the TCGA cohort together with overall survival and metastases-free survival in the natural history cohort. In the TCGA, a high ratio of neoplastic cells in the highly attended patches was significantly associated with shorter progression free survival (p-value=0.01) while high ratios of necrotic, and stromal cells were significantly associated with longer PFS (p-values=0.031 and 0.002, respectively) (Figure 5). Additionally, we found a significant association between the ratio of stromal to neoplastic cells and PFS (p-value=0.006) (Figure 5).

**Figure 5.**
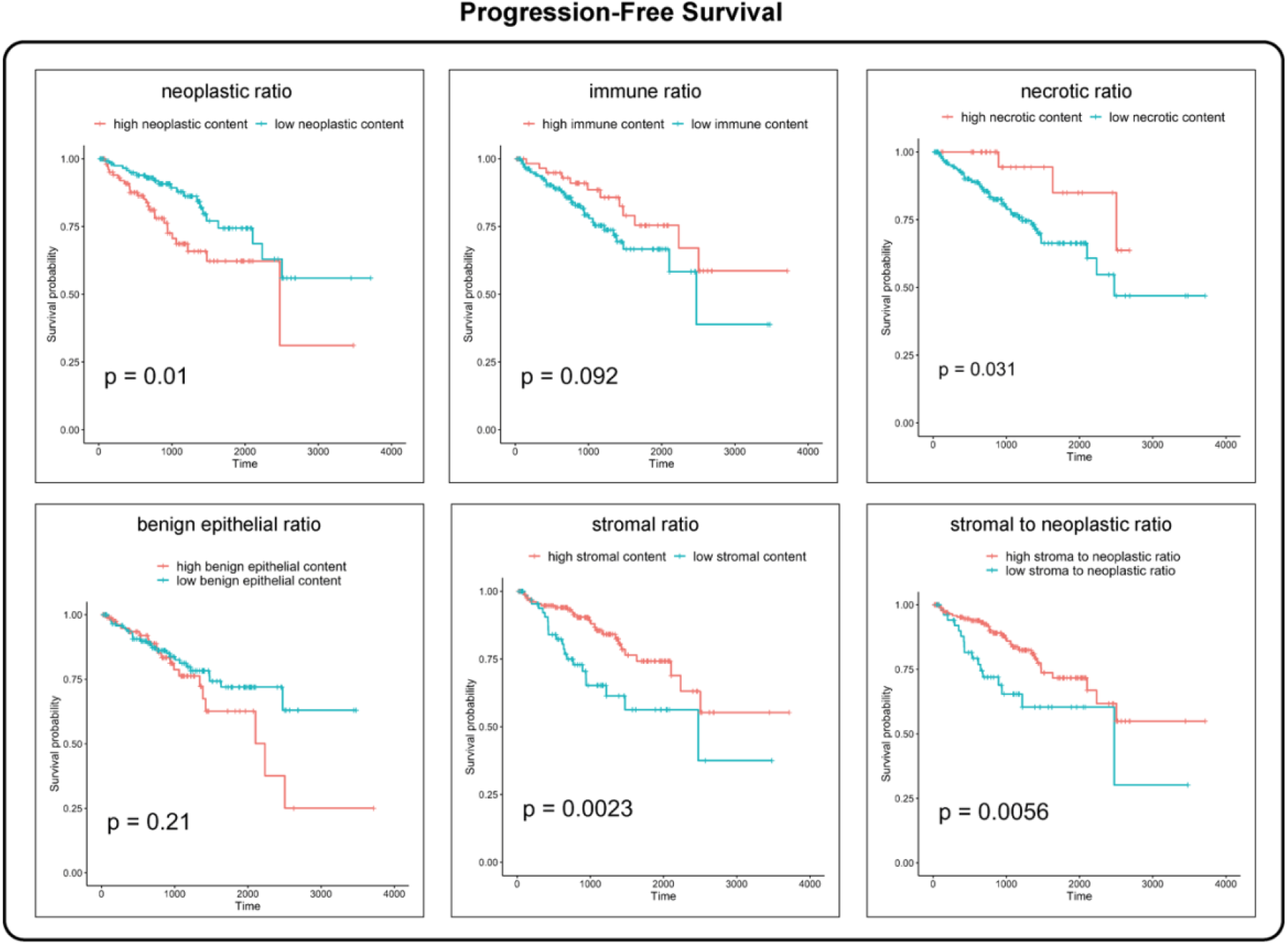
Cellular composition in the highly attended patches is associated with progression-free survival in the TCGA cohort. Kaplan-Meier curves showing the association between the ratio of each nuclear type and progression-free survival. The ratio of each cell type was calculated by dividing the absolute number of that cell type over the number of all cells in the highly attended patches of each slide. The stromal to neoplastic ratio was calculated by dividing the number of stromal cells by that of neoplastic cells in the highly attended patches of each slide. These ratios were then then binarized into high versus low using maximally selected log-rank statistics.

In the natural history cohort, a high ratio of neoplastic cells in the highly attended patches was significantly associated with shorter overall survival (p-value=0.02) while high ratios of immune (p-value=0.004), stromal (p-value=0.002), and stromal to neoplastic cells (p-value=0.01) were significantly associated with longer overall survival (Figure 6A). Similarly, high ratios of immune (p-value=0.01), necrotic (p-value=0.01), benign epithelial (p-value<0.001), stromal (p-value=0.002), and stromal to neoplastic ratio (p-value=0.001) were each associated with significantly longer metastasis-free survival (Figure 6B). Altogether, these results show that the ERG status prediction model is also capable of deciphering gigantic WSIs to capture biologically informative small tissue regions whose cellular composition is associated with survival in PCa patients.

**Figure 6.**
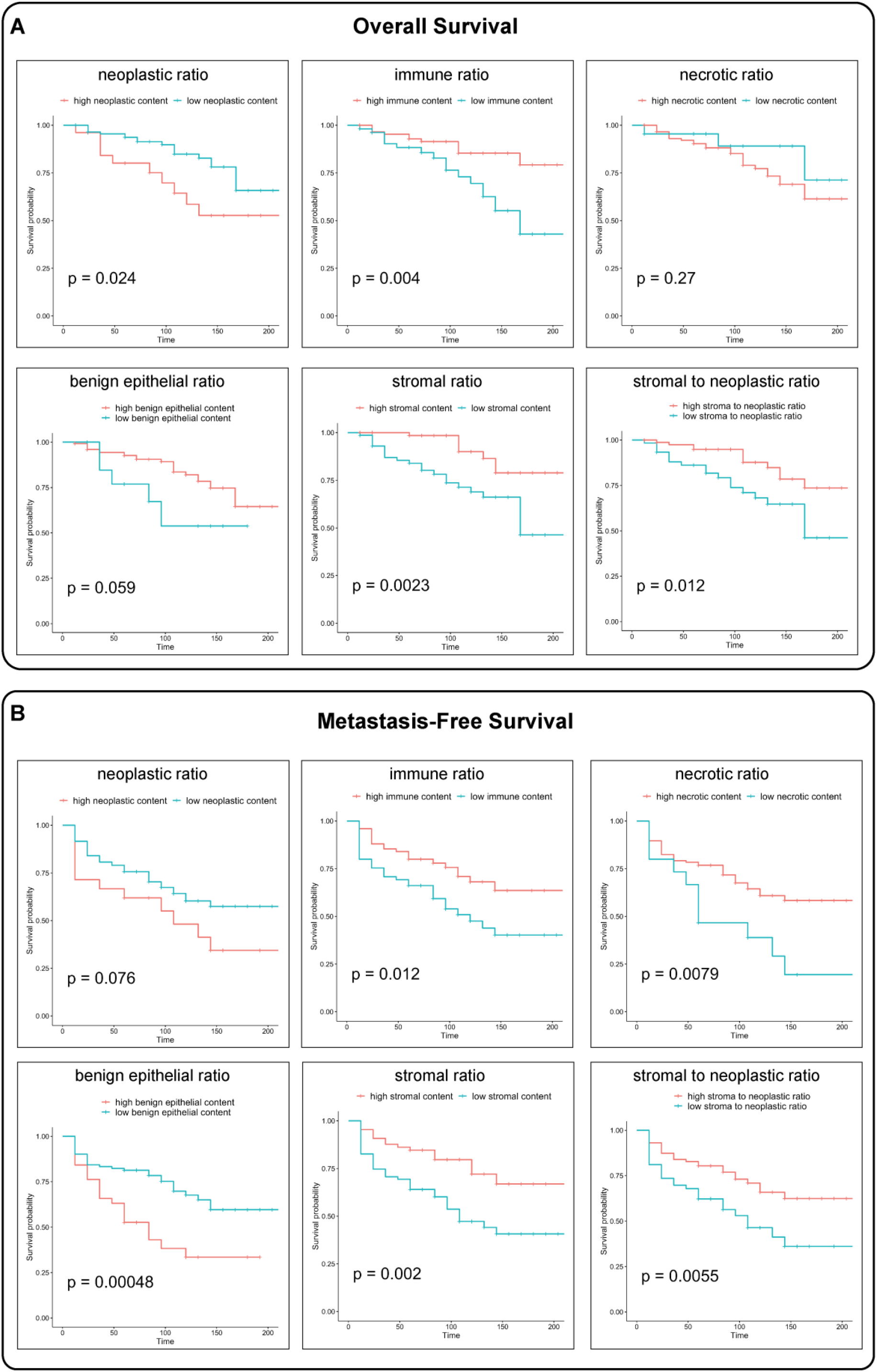
Cellular composition in the highly attended patches is associated with overall and metastasis-free survival in the natural history cohort. **A, B)** Kaplan-Meier curves showing the association between the ratio of each nuclear type and overall survival (A) and metastasis-free survival (B). The ratio of each cell type was calculated by dividing the absolute number of that cell type over the number of all cells in the highly attended patches of each slide. The stromal to neoplastic ratio was calculated by dividing the number of stromal cells by that of neoplastic cells in the highly attended patches of each slide. These ratios were then then binarized into high versus low using maximally selected log-rank statistics.

## Discussion

Over the past few years, there has been a tremendous growth in the research and clinical applications of artificial intelligence (AI) and deep learning (DL) in the fields of computational pathology and cancer research. These applications have allowed for automated or semi-automated inspection of large numbers of histopathology images to extract informative spatially resolved features that can be associated with phenotypes of interest. While deep learning systems have the potential of reducing the workload of pathologists by automatically detecting known morphological features, they can also help detect previously uncharacterized features. Altogether, this has allowed for a wide array of diagnostic and prediction tasks that implement these features to detect certain clinical and molecular phenotypes as well as improve the subtyping of various disease states and cancer types ^12,41^. Particularly, semi-supervised learning has been utilized extensively to implement prediction tasks on WSIs using only slide-level labels instead of pixel-level annotations. This has covered a wide array of research and clinical interests with variable complexity ^42^. For instance, DL systems have been deployed on H&E-stained histopathological slides to detect tumor tissue ^16,17,27^ and for tumor subtyping ^20,22^, grading ^19^, and prognostication ^43–45^. Additionally, DL models have been employed to predict several molecular alterations including for example ER status in breast cancer ^27^, BRAF ^28^ and TP53 ^46^ mutations, and microsatellite instability ^47^. In this study, we introduce a semi-supervised DL model capable of inferring the ERG:TMPRSS2 gene fusion status from digitized H&E-stained WSIs. In the training cohort which included 318 WSIs from the TCGA dataset, the best performing model had an AUC and accuracy of 0.84 and 0.77, respectively and was tested on the TCGA testing set (59 WSIs) with an AUC of 0.72 and accuracy of 0.70. Additionally, we deployed the model on an internal testing cohort (the natural history cohort) with 314 WSIs in which our model maintained its good performance with an AUC of 0.73 and accuracy of 0.69. These results show that our model could maintain its predictive performance on slide cohorts from different institutions and scanned by different technologies.

While many studies in this field have focused on reporting the predictive performance with little regard for biological interpretation, in our study, we thoroughly addressed the interpretability of our model to understand what distinct morphological features are associated with its predictions. Specifically, the use of attention-based DL in our study allowed us to assign attention scores for patches contributing to slide-level representation ^32^ with high scores suggesting the importance of these patches in predicting either ERG-positive or negative cases. With this in mind, we computed the attention scores for all the slide patches and examined the highly attended patches from each slide predicted as either positive or negative to see if there are pathomorphological or cellular composition features specific to ERG status. To characterize the cellular composition in these highly attended patches, we used HoVer-Net model ^35^ trained on the PanNuke dataset ^34^ to segment and classify the nuclei into one of five main nuclear types; neoplastic, immune, stromal (connective tissue), necrotic, and benign epithelium. Notably, highly attended patches for the positive class were enriched in more neoplastic content than their ERG-negative counterparts and were more enriched in necrotic, immune, and stromal cells. We hypothesized that the unique cellular content in these regions might capture prognostic information. For this reason, we examined whether the ratio of each nuclear type is associated with progression-free survival in the TCGA cohort as well as overall survival and metastasis-free survival in the natural history cohort. Notably, the ratio of neoplastic cells in the highly attended patches was significantly associated with shorter progression-free survival and overall survival in the TCGA and natural history cohorts, respectively. In contrast, the ratio of immune cells was associated with longer progression-free (TCGA cohort), overall, and metastasis-free survival (natural history cohort). These results show that the cellular composition in the relevant tissue regions reflects known biology and can serve as additional validation of our results.

In this study, our model could predict the ERG:TMPRSS2 gene fusion status using routine histopathological images and only slide-level labels without expert pixel-level annotation. This together with other studies with similar scope ^28,46,47^ highlight the rising importance and relevance of AI and computer vision in the field of pathology by assisting pathologists and improving the cost and efficiency of the diagnostic process. For instance, detecting ERG:TMPRSS2 fusion is currently performed using either fluorescence in situ hybridization (FISH) or reverse transcription-polymerase chain reaction (RT-PCR), both of which are costly and require trained personnel and equipment. Having clinical-grade models capable of detecting ERG status from simple H&E-stained sections, even on the patient level, can offer tremendous financial and operational advantages. While most of the current models, including ours, did not yet achieve the performance that can enable their clinical usage, they show the potential utility of DL systems to perform complex diagnostic tasks using only tissue pathomorphological features from simple H&E-stained slides without pixel annotations. In fact, it is expected that the accuracy of such models will continue to improve over the next few years with more data available for training and validation up to the point where their clinical deployment will be optimal and justified,

The present study has some inherent limitations. First, our ERG model has been retrospectively tested on two datasets, the first being a subset of the PCa TCGA cohort with 59 slides and the second is the entire natural history cohort with 314 slides. However, there is still the need to prospectively validate the model on larger cohorts from different institutions. Second, while the performance of our model was optimal and stable when tested on slides from a different institution and scanned by a different slide scanner (Hamamatsu versus Aperio), this performance can still improve with further training and re-evaluation on future multi-institutional cohorts. Additionally, our model has been trained and tested on radical prostatectomy specimens and would need additional evaluation on prostate biopsy specimens which would further enhance its clinical utility as a diagnostic or screening test for ERG status. Finally, it is worth mentioning that some deep learning models could achieve a performance similar to or even better than human performance. We believe that with further training and validation on larger scale datasets, we would be able to get higher quality diagnostic networks that can be implemented seamlessly into clinical practice. Perhaps soon, automated ERG fusion prediction from H&E-stained histopathological slides may have the potential of becoming a robust diagnostic and affordable tool that would save the time and efforts associated with costly molecular investigation ^48^.

Here, we presented a DL system that can predict ERG status using digitized H&E-stained WSIs from prostate cancer radical prostatectomy specimens. Such tool can potentially be used by clinicians to infer ERG fusion status quickly and accurately. We thoroughly examined the cellular composition of the highly attended patched for cases predicted as either ERG-positive or negative and found a significant association between this composition and overall, progression-free, and metastasis-free survival. Altogether, these findings show the utility of semi-supervised DL models in predicting a complex phenotype like ERG:TMPRSS2 fusion from routine histopathological slides without known morphological features or pixel-level annotation.

## Declaration of interests

The authors declare no competing interests.

## Author contributions

M.O. and L.M. conceived the research question. M.O., T.L.L., and L.M. collected the datasets. M.O., Z.X., and S.B.R. performed the analysis. M.O. and M.M. wrote the manuscript. M.L., T.L.L., and L.M. supervised the analysis and the manuscript writing. All authors read and approved the final version of the manuscript.

## Data Availability Statement

Digitized whole slide images from the TCGA prostate cancer are publicly available through the GDC data portal and can be accessed using the following link: https://portal.gdc.cancer.gov/. The scripts used to perform this analysis will be made publicly available on GitHub at the time of publication.

## Notes

### Competing Interest Statement

The authors have declared no competing interest.

